# Ex Vivo Removal of CD41 positive platelet microparticles from Plasma by a Medical Device containing a *Galanthus nivalis agglutinin* (GNA) affinity resin

**DOI:** 10.1101/2025.05.09.652772

**Authors:** Rosalia de Necochea Campion, Miguel Pesqueira, Steven P. LaRosa

## Abstract

**Background:** Platelet microparticles (PMP) are elevated in and associated with disease activity in a number of diseases including cancer, neurological conditions, autoimmune diseases, and infectious diseases. This finding raises the possibility of a removal of these microparticles as a therapeutic strategy. The Hemopurifier is an experimental device consisting of a plasma separator and an affinity resin containing *Galanthus nivalis agglutinin* (GNA) affinity resin that has previously been shown to remove extracellular vesicles in vitro and in vivo. In this proof on concept study, we sought to determine ex vivo removal of platelet-derived microparticles from healthy human plasma by the Hemopurifier.

**Methods:** Two hundred milliliters of thawed healthy human plasma were circulated over the Hemopurifier device at a rate of 100 mL/minute. Plasma samples were taken at time points equivalent to a 4-, 6- and 8-hour clinical Hemopurifier session in a healthy adult. Microparticles were isolated from these timepoints and analyzed for treatment concentration changes. Platelet microparticle counts were determined by binding to antiCD41.

**Results:** The Hemopurifier removed 98.5% of platelet microparticles during the equivalent of a 4-hour clinical session.

**Conclusions:** We demonstrated that an extracorporeal device with a GNA affinity resin removes platelet microparticles from normal healthy plasma. Next steps would be demonstration of the removal of PMPs from plasma by the Hemopurifier in different disease states and characterization of the cargo within removed PMPs.

## Introduction

Extracellular vesicles (EVs) are nanoparticles surrounded by a lipid bilayer released by all cell types. These particles are characterized by size with exosomes being 30-100nm in size and of endocytic origin, ectosomes, also known as microparticles or microvesicles, 100-1000 nm in size, and apoptotic cell bodies. Extracellular vesicles contain cargo involved in cell-to-cell communication including proteins, lipids, and nucleic acids ^1^. There is an almost exponential increase in the number of published papers in the peer-reviewed literature describing the role of EVs in different disease states.

The vast majority of extracellular vesicles found in the circulation are derived from platelets (PD-EVs). Estimates are that Platelet-derived EVs comprise up to 70-90% of the total EV population as determined by the surface marker CD41 ^1^. Platelet derived EVs have been determined to have effects on the endothelium, immune cells and tumor cells. Elevated numbers of large CD41+ve extracellular vesicles (platelet microparticles (PMPs)) have been implicated in cancers, as well as autoimmune, neurologic, and cardiovascular diseases ^2^. This finding raises the hypothesis that removing platelet-derived microparticles might improve these conditions.

An investigational extracorporeal device called the Hemopurifier® (HP) (Aethlon Medial, Inc., San Diego, CA, USA) has been developed to bind and remove extracellular vesicles. The device is a single-use plasma separator cartridge that is modified to contain a proprietary affinity resin in the plasma space externa; composed of the lectin external to hollow fibers with a nominal pore size of 200-500nm ^3^. The affinity resin contains the plant lectin *Galanthus nivalis* agglutinin (GNA) covalently bound to a medical grade diatomaceous earth. GNA has been found to bind to terminal non-reducing or reducing mannose as well as alpha-1,3 linked mannose oligosaccharides which have been found on the surface of extracellular vesicles ^4,5^. Both *in-vitro* and *in-vivo* binding of EVs by the Aethlon HP has been demonstrated ^6,7,8^.

Specific binding of platelet microparticles by the Hemopurifier has not been previously examined. We performed a proof-of-concept ex vivo experiment utilizing donor normal healthy plasma. Briefly, the plasma was thawed and pumped over a Hemopurifier to simulate the duration of a clinical trial Hemopurifier session. Serial samples were taken to examine the number of CD41+ve EVs remaining at given time points.

## MATERIALS AND METHODS

### Hemopurifier Treatment of Pooled Human Plasma

De-identified, apheresis-derived pooled normal human plasma was obtained from an FDA-approved collection center (Innovative Research, Novi, MI, USA) using K□EDTA as the anticoagulant. The plasma tested negative for a panel of viral markers (Innovative Research; catalog #IPLAK2E1000ML, Lot#43968). According to the supplier, all plasma was collected in compliance with FDA regulations. Samples were fully de-identified prior to purchase, and no identifiable donor information was made available to the researchers.

A total of 300 mL of pooled human plasma was centrifuged at 1,000 × *g* for 5 minutes to remove large cellular debris. Following clarification, 250 mL of the supernatant was transferred to a reservoir for HP treatment circulation, then 50 mL of plasma was collected and stored to serve as the pre-treatment (baseline) control. Prior to initiating treatment of the remaining 200 mL of plasma, the HP circuit was flushed with 1 L of sterile 0.9% saline to ensure proper equilibration. The 200 mL of plasma was then circulated through the Hemopurifier at a flow rate of 100 mL/min, and plasma samples were collected after 9.6, 14.4, and 19.6 total volume passes. This sample schedule corresponded to HP clinical treatment durations equivalent to 4-, 6-, and 8-hours treatment, when considering that an average adult human has a 5L blood volume and the HP clinical procedure can be performed using an extracorporeal flow rate of 200 mL/min. In this experiment, baseline and HP treated plasma samples were stored frozen in 1 mL aliquots at −80°C.

### Extracellular Vesicle Isolation

Microparticles (extracellular vesicles with a size of 100-500 nm) were isolated from 1 mL of baseline and timepoint plasma samples by first performing two pre-clearing centrifugation steps of 2,500 x g for 15 minutes at room temperature, recovering 850 µL of supernatant and further centrifugating at 18,000 x g for 45 minutes at 4°C to pellet the EVs. Plasma supernatant was completely removed, and large EV pellets were resuspended in 100 µL of 0.22 µm PES membrane-filtered PBS, incubated for 45 minutes at room temperature to allow for complete resuspension, then separated into 25 µl aliquots and stored at −80°C prior to analysis.

### Nanoparticle Flow Cytometry Analysis

Extracellular vesicles were quantified using the single-particle detection NanoFCM™ Flow Analyzer U30 instrument (NanoFCM, Inc., Xiamen, China). The total EVs were measured using the phospholipid bilayer dye MemGlow488nm (MemGlow™ 488: Fluorogenic Membrane Probe, Cytoskeleton). The concentration and size distribution profile of the EVs was determined utilizing the manufacturer’s provided calibration silica beads kit (NanoFCM™ Silica Nanospheres Cocktail #3; catalog # S17M-MV). The identification of platelet microvesicles was determined through the staining, and a one-hour incubation at room temperature, of 100 nM MemGlow488nm and APC anti-CD41 (BioLegend Inc., San Diego, CA, USA; catalog #303704).

## RESULTS

As shown in Figure 1A, we observed the total number of 100-500 nm microparticles measured at baseline to be just over 4E+08/mL of plasma with more than 96% of these microparticles removed after a 4-hour Hemopurifier treatment, and not much additional depletion at subsequent time intervals. This indicates that the Aethlon Hemopurifier is highly effective in depleting microparticles in this size range and the 4-hour treatment appears sufficient for maximum removal. The total number of CD41+ve microparticles at baseline was 1.93E+07/mL of plasma with over 98.5% of these microparticles removed after a 4-hour treatment with the Hemopurifier and subsequently undetectable at later time points (Figure 1B).

**Figure 1.**
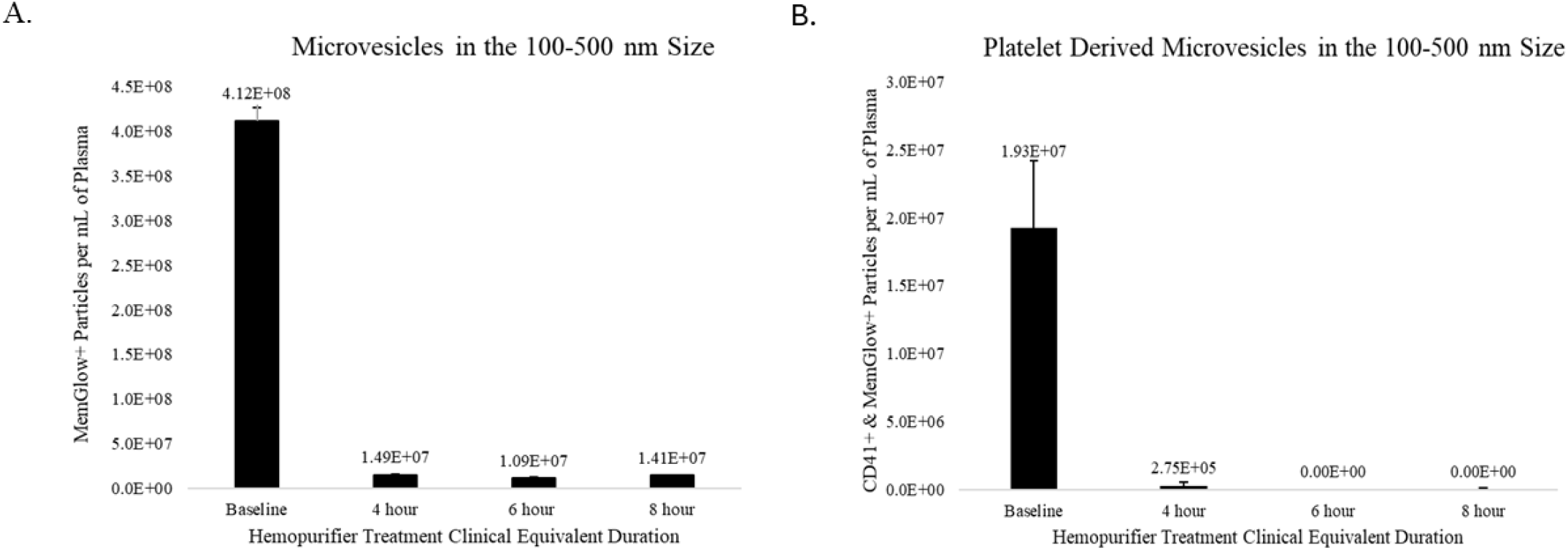
Depletion of plasma microvesicles during Hemopurifier Treatment. Nanoparticle flow cytometry analysis of microparticles isolated from pooled human plasma HP treatment timepoint samples demonstrate a concentration reduction of varied microparticle populations in the 100-500 nm size range: (A) MemGlow™ 488+ microparticles, otherwise referred to as microvesicles, and (B) platelet-derived microvesicles co-localized with antiCD41-APC and MemGlow™ 488. The data shown above represents the average of two technical replicates and their respective standard deviation. Limit of detection was 5.78E+04 particles/mL of plasma.

## DISCUSSION

In this proof on concept ex vivo study, we show for the first time that an extracorporeal medical device that includes an affinity resin containing the plant lectin *Galanthus nivalis agglutinin* removes the vast majority of platelet microparticles from a large volume of healthy human plasma during a 4–6-hour treatment period. The data from this study builds on the previously described in vitro and in vivo binding of extracellular vesicles by the Hemopurifier device. In one in vitro experiment 92-99% of EVs isolated from patients with distinct tumor types including head and neck cancer, melanoma, ovarian cancer, esophageal cancer, and breast cancer were removed by a bench top Hemopurifier when these EVs were suspended in buffer ^6^. In the second experiment, between 31-62% of EVs were removed directly from a plasma sample of a patient with non-small cell lung cancer by the bench top Hemopurifier ^7^. *In vivo* reduction of EVs was measured during a total of ten HP sessions in 2 patients. In a patient with severe COVID-19 infection and multi-organ system failure a downward trajectory in total exosome concentration was observed over the course of the eight treatments ^8^. In a patient with head and neck cancer EV counts decreased after 2 treatments 21 days apart with a slow rise to levels below the baseline measurement at 7 days post treatment ^7^.

The Hemopurifier is currently being studied in an Australian clinical trial in patients with solid tumors who are not responding to anti PD-1 therapy. Platelet microparticles have previously been demonstrated to play a role in the progression of cancer by promoting tissue invasion, metastases, angiogenesis, and inhibition of apoptosis ^1^. Elevated platelet microparticles have been found in patients with non-small lung cancer progressing during treatment with the Nivolumab and Pembrolizumab. A platelet microparticle count of > 80,000 events/ml had an ROC AUC of 0.825 for progression of disease during therapy ^9^. Another study by Wang et al found a platelet derived activated extracellular vesicle count of 62,700/ml at 3 months into therapy was predictive of outcome in this population ^10^. As part of this ongoing clinical trial, participants will have plasma samples taken before, during, and once weekly for 8 weeks post-Hemopurifier treatment for measurements of extracellular vesicles ^7^. Platelet microparticles (PMPs) will be a population of EVs specifically evaluated pre and post Hemopurifier treatment.

Platelet Microparticles as a therapeutic target for intervention extends beyond oncologic indications. In the neurological space, PMPs have been studied in Alzheimer’s disease (AD) and Multiple Sclerosis (MS). Odaka et al compared sera between healthy donors and patients with Alzheimer’s disease ^11^. The extracellular vesicles from AD patients had significantly greater staining for the PD EV marker CD41. Marcos-Ramiro and colleagues compared plasma from patients with all forms of multiple sclerosis (clinically isolated syndrome, Relapsing Remitting MS, secondary progressive MS, and primary progressive MS) to healthy controls ^12^Analysis of the pool of all MS patients revealed that the platelet microparticles counts were statistically significantly greater in the MS population than in the healthy controls (27,203 counts/microliter+/− 16.767 vs. 15,646 counts/microliter+/− 11,901; p<0.001). Upon subgroup analysis this statistically significant difference held for all forms of MS except for clinically isolated syndrome. Additionally, these authors demonstrated that these platelet microparticles from MS patients disturbed endothelial barrier function.

Platelet microparticles have also been implicated in the pathogenesis of autoimmune diseases. In Rheumatoid Arthritis, the levels of circulating platelet microparticles were greater in patients with active disease manifestations and elevated acute phase reactant levels than in patients with quiescent disease ^13^. Consumption of large platelets with production of platelet microparticles has also been described in systemic lupus erythematosus (SLE) ^14^. Levels of platelet microparticles were 2-7 X higher in patients with SLE than healthy controls. Microparticles in patients were associated with coagulation and inflammatory markers as well as impaired renal function. Finally, platelet microparticles composed 67% pf microparticles found in patients with systemic sclerosis and were greater in patients than healthy controls (58.8 × 10^5^ per ml vs 30.4 × 10^5^/ml; p<0.05) ^15^. Platelet microparticles in systemic sclerosis contribute to the pathology seen in part through neutrophil activation and neutrophil extracellular trap (NET) formation ^16^.

An additional disease area where platelet microparticles have been studied is in infectious diseases. In sepsis, a dysregulated response to infection resulting in organ dysfunction, levels of platelet microparticles were significantly greater in patients with sepsis than healthy controls, and platelet microparticles correlated with components of neutrophil extracellular traps. Infusion of platelet microparticles in a mouse sepsis model induced NET formation and endothelial dysfunction ^17^. Platelet microparticle counts are associated with increased disease severity in COVID-19 potentially mediated through HMGB-1 ^18^. Elevated platelet extracellular vesicle levels have also been observed in patients with persistent symptoms following acute COVID-19 infection ^19^. In Dengue fever, PMPs are associated with thrombocytopenia and vascular leakage ^20^. Platelet microparticles can participate in the pathogenesis of cerebral malaria through activation of clotting and binding to the brain endothelium ^21^.

This initial pre-clinical proof of concept study has limitations. Healthy human plasma was used to assess the removal of platelet microparticles by the Hemopurifier. It is possible that PMPs from different disease states may have differential binding to the device. Future studies will need to be done on plasma from disease states of interest. In this study we did not assess the changes in cargo components of the platelet microparticles removed. In the future we will need to determine that removal of platelet microparticles translates into removal of cargo that are known mediators of inflammation, coagulation, or cancer progression. Finally, we did not perform a lectin microarray on the platelet microparticles to confirm specific binding to the GNA lectin. However, lectin microarray analysis performed on platelet microparticles from Alzheimer’s disease patients bound to GNA, suggests that the mannose target for this lectin is on Platelet-derived EVs^11^.

In conclusion, the Hemopurifier device was able to remove platelet microparticles from healthy normal plasma. An ongoing clinical trial in patients with solid tumors will yield data on the removal of platelet microparticles in vivo. Additional ex vivo platelet microparticle removal studies will be performed in non-oncologic indications.

## Notes

**Conflict of interests** Authors SPL, MP, and RdNC are employees of Aethlon Medical, Inc.

### Competing Interest Statement

Authors SPL, MP, and RdNC are employees of Aethlon Medical, Inc. which funded the study.

## REFERENCES

1. Żmigrodzka M, Guzera M, Miśkiewicz A, et al. The biology of extracellular vesicles with focus on platelet microparticles and their role in cancer development and progression. Tumour Biol. 2016;37(11):14391–14401. doi: 10.1007/s13277-016-5358-6.

2. Eustes AS, Dayal S. The Role of Platelet-Derived Extracellular Vesicles in Immune-Mediated Thrombosis. Int J Mol Sci. 2022;23(14) :7837. doi:10.3390/ijms23147837.

3. Marleau AM, Chen CS, Joyce JA, Tullis RH. Exosome removal as a therapeutic adjuvant in cancer. J Transl Med. 2012;10 :134. doi:10.1186/1479-5876-10-134.

4. Hester G, Wright CS. The Mannose-specific Bulb Lectin from Galanthus nivalis (Snowdrop) Binds Mono- and Dimannosides at Distinct Sites. Structure Analysis of Refined Complexes at 2.3 Å and 3.0 Å Resolution. J. Mol. Biol. 1996; 262: 516–531.

5. Batista BS, Eng WS, Pilobello KT, et al. Identification of a conserved glycan signature for microvesicles. J Proteome Res. 2011; Oct 7:10(10):4624–33. doi: 10.1021/pr200434y.

6. Marleau AM, Jacobs MT, Gruber N, et al. Abstract 4509: Targeting tumor-derived exosomes using a lectin affinity hemofiltration device. Cancer Res 2020;80 (16_Supplement):4509.

7. Brown MP, Matos M, Clarke S, et al. Protocol for a Safety, Feasibility, and Dose-Finding Study of the Hemopurifier® Device in Patients with Solid Tumors Who Have Stable or Progressive Disease During Pembrolizumab or Nivolumab Monotherapy. medRxiv [Preprint] 2025. DOI: 10.1101/2025.03.20.25323761.

8. Amundson, DE, Shah US, de Necochea-Campion R, et al. Removal of COVID-19 Spike Protein, Whole Virus, Exosomes, and Exosomal MicroRNAs by the Hemopurifier® Lectin-Affinity Cartridge in Critically Ill Patients With COVID-19 Infection. Front Med (Lausanne) 2021; 8: 744141.doi: 10.3389/fmed.2021.744141. eCollection 2021.

9. Liu T, Wang J, Li T, et al. Predicting disease progression in advanced non-small cell lung cancer with circulating neutrophil-derived and platelet-derived microparticles. BMC Cancer. 2021;21(1):939. doi: 10.1186/s12885-021-08628-4.

10. Wang CC, Tseng CC, Chang HC, et al. Circulating microparticles are prognostic biomarkers in advanced non-small cell lung cancer patients. Oncotarget. 2017;8(44):75952–75967. doi: 10.18632/oncotarget.18372

11. Odaka H, Hiemori K, Shimoda A, et al. Platelet-derived extracellular vesicles are increased in sera of Alzheimer’s disease patients, as revealed by Tim4-based assays. FEBS Open Bio. 2021;11(3):741–752. doi: 10.1002/2211-5463.13068.

12. Marcos-Ramiro B, Oliva Nacarino P, Serrano-Pertierra E, et al. Microparticles in multiple sclerosis and clinically isolated syndrome: effect on endothelial barrier function. BMC Neurosci. 2014; 15:110. doi: 10.1186/1471-2202-15-110.

13. Knijff-Dutmer EA, Koerts J, Nieuwland R, et al. Elevated levels of platelet microparticles are associated with disease activity in rheumatoid arthritis. Arthritis Rheum. 2002;46(6):1498–503. doi: 10.1002/art.10312.

14. Mobarrez F, Fuzzi E, Gunnarsson I, et al. Microparticles in the blood of patients with SLE: Size, content of mitochondria and role in circulating immune complexes. J Autoimmun. 2019; 102:142–149. doi: 10.1016/j.jaut.2019.05.003.

15. Guiducci S, Distler JH, Jüngel A, et al. The relationship between plasma microparticles and disease manifestations in patients with systemic sclerosis. Arthritis Rheum. 2008;58(9):2845–53. doi: 10.1002/art.23735.

16. Maugeri N, Campana L, Gavina M, et al. Activated platelets present high mobility group box 1 to neutrophils, inducing autophagy and promoting the extrusion of neutrophil extracellular traps. J Thromb Haemost. 2014;12(12):2074–88. doi: 10.1111/jth.12710.

17. Jiang M, Wu W, Xia Y, et al. Platelet-derived extracellular vesicles promote endothelial dysfunction in sepsis by enhancing neutrophil extracellular traps. BMC Immunol. 2023;24(1):22. doi: 10.1186/s12865-023-00560-5.

18. Maugeri N, De Lorenzo R, Clementi N, et al. Unconventional CD147-dependent platelet activation elicited by SARS-CoV-2 in COVID-19. J Thromb Haemost. 2022;20(2):434–448. doi: 10.1111/jth.15575.

19. Campello E, Radu CM, Simion C, et al. Longitudinal Trend of Plasma Concentrations of Extracellular Vesicles in Patients Hospitalized for COVID-19. Front Cell Dev Biol. 2022; 9:770463. doi: 10.3389/fcell.2021.770463.

20. Patil R, Bajpai S, Ghosh K, Shetty S. Microparticles as prognostic biomarkers in dengue virus infection. Acta Trop. 2018; 181:21–24. doi: 10.1016/j.actatropica.2018.01.017.

21. Pankoui Mfonkeu JB, Gouado I, Fotso Kuaté H, et al. Elevated cell-specific microparticles are a biological marker for cerebral dysfunctions in human severe malaria. PLoS One. 2010 14;5(10): e13415. doi: 10.1371/journal.pone.0013415.

